# Phenotype Prediction Using BEN-Based Predictive Modeling (BPM)

**DOI:** 10.1101/2024.09.07.611838

**Authors:** Donghui Song, Ze Wang

**Author notes:** Corresponding author: Donghui Song, No.19, Xinjiekouwai St, Haidian District, Beijing, 100875, CN, Corresponding author: Ze Wang, Ph.D., 670 W Baltimore St, Baltimore, MD, 21201, USA.

## Abstract

Functional connectivity (FC) has been successfully used to predict cognitive functions, behaviors, and other phenotypes using connectome-based predictive modeling (CPM). One limitation is that FC reflects the covariation or synchrony relationship of the temporal profile in two regions and neglects the local temporal features of the brain activity. In this study, we used brain entropy (BEN) mapping to characterize regional brain activity and developed a BEN-Based Predictive Modeling (BPM) to predict phenotype. BEN measures the disorder and complexity of brain activity and has been shown to effectively capture brain activity features related to cognition and neurological disorders. Using data from the HCP 7T fMRI and corresponding 3T structural images, we calculated gray matter volume (GMV) as well as BEN from resting state and movie-watching data. We constructed prediction models based on BEN and GMV using different numbers of parcellation of the brain atlas, applying 10-fold cross-validation. Our results indicated that the BEN-based predictive model not only outperformed GMV-based predictive modeling but also achieved prediction accuracy comparable to that of CPM. Our study demonstrates that BEN can capture extensive brain activity information for accurate phenotype prediction, providing information at least as valuable as that from connectome. Additionally, our research lays the groundwork for future applications of BPM in developmental and clinical practice.

## 1. Introduction

Neuroimaging has been widely used to draw populational knowledge about brain structures and functions as well as phenotypes. Individual differences of neuroimaging data which are ignored in the populational analysis are actually equally important as a crucial goal of neuroimaging is to use the individual brain measures to predict cognition, behavior, and others phenotype at the individual level. Over the past decade, the connectome-based predictive modeling (CPM) proposed by Emily and Shen et al (Finn, Shen et al. 2015, Shen, Finn et al. 2017) has gain increasing popularity in personalized phenotype prediction. The successful applications include intelligence (Anderson and Barbey 2023, Wilcox and Barbey 2023), emotion (Ju, Horien et al. 2020, Wang, Goerlich et al. 2021), and addiction (Yip, Scheinost et al. 2019, Lichenstein, Scheinost et al. 2021) etc. Despite the growing emphasis on network-based analyses of brain function, local temporal dynamics are still crucial and are not fully captured by functional connectivity (FC). Based on the inter-regional correlation, CPM basically ignores properties of local temporal dynamics, which certainly represents a mis-opportunity. Therefore, there is a pressing need to develop a straightforward and practical predictive model based on local temporal dynamic features.

Brain entropy (BEN) quantifies the nonlinear dynamical characteristics of brain activity, representing the irregularity, disorder, or complexity of brain functions (Wang, Li et al. 2014). Our proposed fMRI-based BEN using sample entropy effectively reflects normal brain function and disease states, and it has become an increasingly popular approach for analyzing localized brain activity (Wang, Li et al. 2014, Liu, Song et al. 2020, Wang 2021, Camargo, Del Mauro et al. 2024, Dong-Hui Song 2024). In our previous studies, we characterized the spatial distribution patterns of brain entropy in healthy subjects using large cohort data, and these distribution patterns align with some functional brain networks such as the motor cortex, default mode network, and visual network (Wang, Li et al. 2014). Moreover, our demonstrated that BEN is associated with cognitive functions (Wang 2021, Del Mauro and Wang 2024) and task performances (Lin, Chang et al. 2022, Camargo, Del Mauro et al. 2024, Song and Wang 2024). Importantly, BEN can reflect changes in neuromodulation (Song, Chang et al. 2019, Liu, Song et al. 2024, Song, Deng et al. 2024), pharmacology (Chang, Song et al. 2018, Liu, Song et al. 2020), neurochemical signals (Song and Wang 2024), and disease-related brain activity (Xue, Yu et al. 2019, Wang and Initiative 2020, Jiang, Cai et al. 2023, Del Mauro, Sevel et al. 2024, Dong-Hui Song 2024). These studies strongly suggest that BEN effectively captures the dynamic characteristics of local neural activity under various conditions, demonstrating sensitivity to changes in neural activity. However, for a long time, the relationship between BEN and phenotypes has often been assessed through brain-wide association studies (BWAS), which typically require a large sample size for normal adults (Marek, Tervo-Clemmens et al. 2022, Liu, Abdellaoui et al. 2023). There has been a lack of studies truly predicting phenotypes based on BEN, which limits its application in individualization. In the study, we aim to evaluate whether BEN can effectively predict phenotypes and assess its predictive efficacy.

Gray matter volume (GMV) from voxel-based morphometry (Ashburner and Friston 2000, Mechelli, Price et al. 2005) in vivo using structural MRI (sMRI) also was recently used to predict phenotypes, such as reading comprehension ability (Cui, Su et al. 2018) and intelligence (Hilger, Winter et al. 2020), achieving good predictive performance. However, a challenge is that structural information often lacks functional characteristics, such predictions may demonstrate good performance for the most stable phenotypes but lower performance for more variable phenotypes along function. Our recent study also indicates that resting BEN shows a brain-wise negative correlation with GMV in the lateral frontal and temporal lobes, inferior parietal lobules, and precuneus (Del Mauro and Wang 2024). In this study, we also use GMV-based predictive modeling (GPM) to predict phenotypes and directly compare with BEN-based predictive modeling (BPM). Recently Finn et al also suggested that CPM using movie-watching outperforms rest for the prediction of behavior using CPM (Finn and Bandettini 2021). Furthermore, our study also indicated that BEN while watching movies has higher test-retest reliability compared to resting, and BEN can capture movie-related brain activities (Song and Wang 2024). Therefore, in this study, we will also assess whether BEN during movie-watching provides better phenotype prediction compared to resting BEN.

## 2. Methods

### 2.1 Dataset

All participants from the Human Connectome Project (HCP) 7T release. One hundred seventy-six (106 female) out of 184 participants completed T1w, four resting state (REST) runs, and four movie-watching (MOVIE) runs over two days. All participants were healthy individuals between 22 and 36 years (mean age = 29.4, standard deviation = 3.3).

The structural images were scanned on a 3T Siemens Skyra scanner using a standard 32-channel head coil at Washington University. The functional images were acquired on a 7 Tesla Siemens Magnetom scanner at the Center for Magnetic Resonance Research at the University of Minnesota. Detailed information about data acquisition and preprocessing can be found in (Glasser, Sotiropoulos et al. 2013, Van Essen, Smith et al. 2013), also see (Song and Wang 2024).

### 2.2 GMV

The structural images were preprocessed including distortion correction and bias field correction by HCP named “T1w_acpc_dc_restore_brain.nii.gz”. GMV was computed by the FSL pipeline (Smith, Jenkinson et al. 2004, Douaud, Smith et al. 2007) following FSLVBM protocol (https://fsl.fmrib.ox.ac.uk/fsl/fslwiki/FSLVBM), fslvbm_2_template was used to create the study-specific GM template, fslvbm_3_proc was used to register all GM images to the study-specific template non-linearly. The detailed guide of FSLVBM is available at https://fsl.fmrib.ox.ac.uk/fsl/fslwiki/FSLVBM/UserGuide.

### 2.3 BEN mapping

BEN maps were calculated from our previous study (Song and Wang 2024).

### 2.4 Behavioral data

The HCP database contains a large number of phenotypic measurements. Following the study from Emily S. Finn and Peter A. Bandettini (Finn and Bandettini 2021), we applied these reduced-dimensional data by performing principal components analysis (PCA), which resulted in two primary domains: cognition, and emotion. These reduced-dimensional data are available at https://github.com/esfinn/movie_cpm/tree/master/data. For details on the dimensionality reduction methods used, please refer to (Finn and Bandettini 2021).

### 2.6 Brain Atlas

We used schaefer_2018 (Schaefer, Kong et al. 2018) to extract the mean BEN from each parcel. This atlas provides a range of parcels from 100 to 1000, we selected several parcels from 200 to 600, which helps us determine the optimal number of parcels for generating more stable and accurate predictions.

### 2.7 BEN and GMV-based predictive modeling

We employed a linear regression model from scikit-learn (Pedregosa, Varoquaux et al. 2011). The steps involved in BEN and GMV-based predictive modeling are as follows:

1. First, extract the average BEN values or GMV values for each ROI for all subjects.
2. Given the average BEN values or GMV values of ROI and behavioral scores for all subjects, divide the data into training and testing sets.
3. Build a linear model relating the average BEN or GMV values (x) of each parcel to behavior score (y):

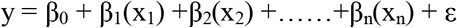

where n is the number of parcels,
4. Calculate average BEN or GMV values (x) for each subject in the test set and use these as input to the linear model in 3 to generate predicted behavior scores (y).

Cross-validation (CV) was conducted using a 10-fold CV. Prediction accuracy was assessed by Person’s coefficient between predicted (model-generated) and observed (true) scores across subjects. To assess the statistical significance of prediction accuracies, we performed a permutation test by shuffling behavior scores for BEN or GMV values and reperforming the entire analysis pipeline to generate a null distribution of expected accuracies due to chance. A total of 10,00 randomizations were performed, then calculated a non-parametric p-value by the following formula,

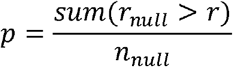

where *n*_*null*_ = 10,00 and r is the correlation coefficient from true models, is the correlation coefficient from the permutation test.

To visualize the prediction weights β for each parcel, we averaged the weights from the CV and then mapped these averaged weights onto the whole brain.

## 3. Results

Cognition scores could be predicted by rsBEN with significant accuracy across 200 parcels to 600 parcels (200 parcels: r=0.130, p=0.046,300 parcels: r=0.248, p=0.003, 400 parcels: r=0.259, p=0.003, 500 parcels: r=0.318, p<0.001, 600 parcels: r=0.275, p=0.001, Fig.1). Cognition scores could be predicted by mvBEN with significant accuracy across 200 parcels to 600 parcels (200 parcels: r=0.163, p=0.003, 300 parcels: r=0.430, p<0.001, 400 parcels: r=0.392, p<0.001, 500 parcels: r=0.348, p<0.001, 600 parcels: r=0.243, p=0.001, Fig.1). Cognition scores could be predicted by GMV with significant accuracy across 300 parcels to 600 parcels (200 parcels: r=0.093, p=0.087, 300 parcels: r=0.175, p=0.004, 400 parcels: r=0.199, p=0.004, 500 parcels: r=0.168, p=0.011, 600 parcels: r=0.263, p<0.001 Fig.1).

**Fig1.**
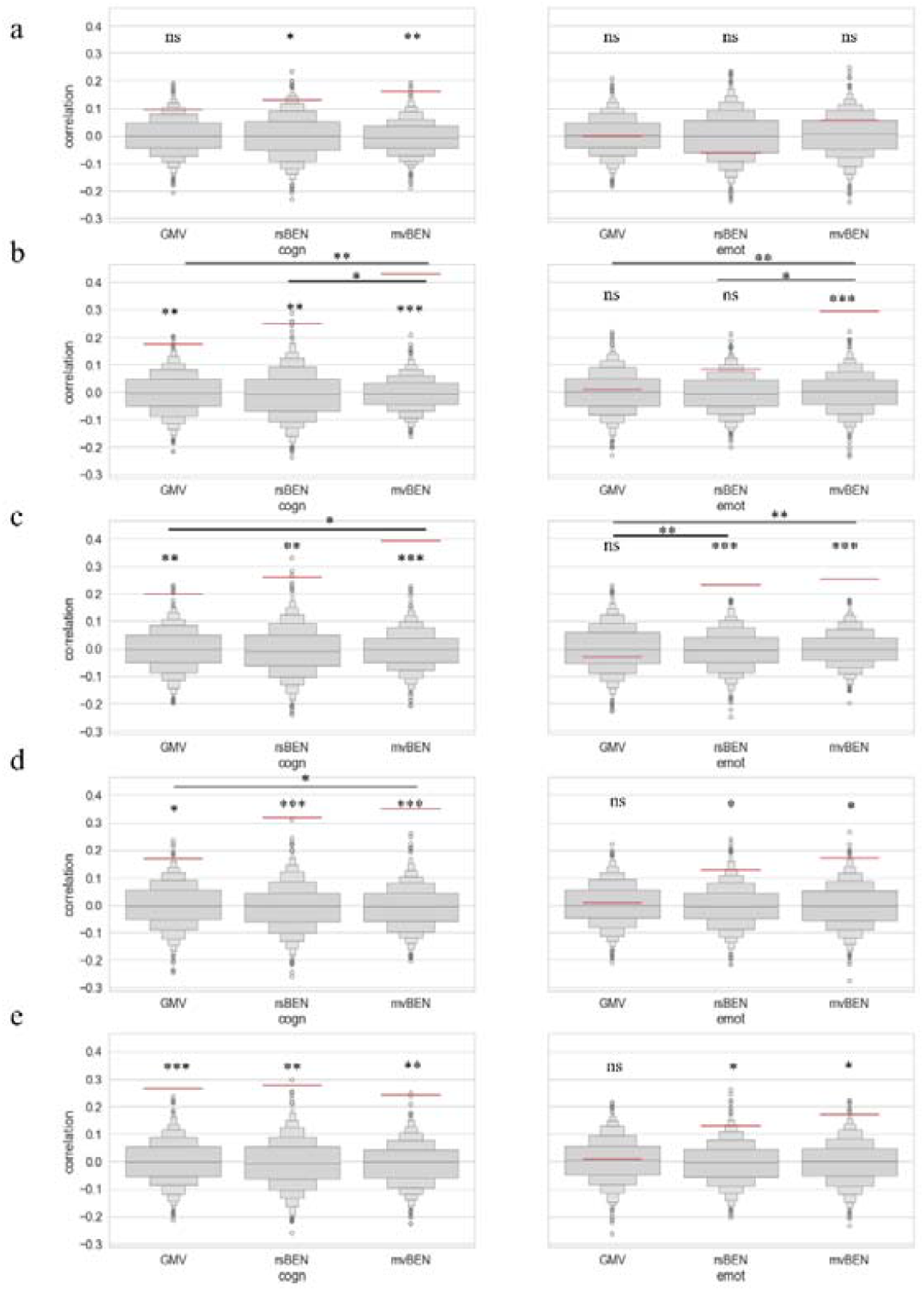
Prediction accuracy. a. 200 parcels, b. 300 parcels, c. 400 parcels, d. 500 parcels, e. 600 parcels. Left is the accuracy for the cognition (cogn) score, and right is the accuracy for the emotion (emot) score. Accuracy was measured as the Pearson’s correlation between predicted and observed scores (y-axis). Light gray boxen plots show null distribution from 10,00 permutations in which behavior scores and GMV, rsBEN, and mvBEN were randomized across subjects. The red horizontal line denotes accuracy for true models. n.s., not significant (*p* > 0.05); \**p* < 0.05; \*\**p* < 0.01; \*\*\**p* < 0.001.

In terms of absolute values, except for the 600 parcels, both rsBEN and mvBEN demonstrate better predictive ability for cognitive scores compared to GMV in all other parcel numbers. Specifically, for parcel numbers ranging from 300 to 500, mvBEN showed significantly better predictive ability for cognitive scores compared to GMV (300 parcels: p=0.004,400 parcels: p=0.024, 500 parcels: p=0.036, Fig.1). Furthermore, at 300 parcels, mvBEN predictive ability for cognitive scores is significantly superior to that of rsBEN (p=0.027, Fig.1).

Emotion scores could be predicted by rsBEN with significant accuracy across 400 parcels to 600 parcels (200 parcels: r=-0.063, p=0.763,300 parcels: r=0.083, p=0.101, 400 parcels: r=0.230, p<0.001, 500 parcels: r=0.128, p<0.032, 600 parcels: r=0.143, p=0.019, Fig.1). Emotion scores could be predicted by mvBEN with significant accuracy across 300 parcels to 600 parcels (200 parcels: r=0.057, p=0.257, 300 parcels: r=0.293, p<0.001, 400 parcels: r=0.253, p<0.001, 500 parcels: r=0.171, p=0.016, 600 parcels: r=0.145, p=0.025, Fig.1). Emotion scores couldn’t be predicted by GMV with significant accuracy at any parcels (200 parcels: r=-0.001, p=0.526, 300 parcels: r=0.008, p=0.459, 400 parcels: r=-0.033, p=0.651, 500 parcels: r=0.007, p=0.465, 600 parcels: r=-0.02, p=0.625, Fig.1).

In terms of absolute values, both rsBEN and mvBEN demonstrate a better predictive ability for emotion scores compared to GMV across 300 parcels to 600 parcels. Specifically, for parcel numbers ranging from 300 to 400 parcels, mvBEN showed significantly better predictive ability for emotion scores compared to GMV (300 parcels: p=0.003,400 parcels: p=0.003, Fig.1). Furthermore, at 300 parcels, the predictive ability of mvBEN for emotion scores is significantly superior to that of rsBEN (p=0.021, Fig.1), at 400 parcels, the predictive ability of rsBEN for emotion scores is significantly superior to that of GMV (p=0.006, Fig.1).

We next examined which parcels were most important to the predictive models for cognitive scores and emotion scores. We average the β values for each parcel of the predictive model from the training set and then project them onto the whole brain (Fig2, Fig3). The results indicated that predictive capability is associated with a broad range of brain regions. We focus on the weights exhibited by the parcels with the highest predictive power. Specifically, for the GMV-based prediction model of cognitive scores, we selected 600 parcels. For the rsBEN-based prediction model of cognitive scores, we chose 500 parcels. For the mvBEN-based prediction model of cognitive scores, we selected 300 parcels. For the rsBEN-based prediction model of emotion scores, we opted for 400 parcels, and for the mvBEN-based prediction model of emotion scores, we chose 300 parcels.

**Fig2.**
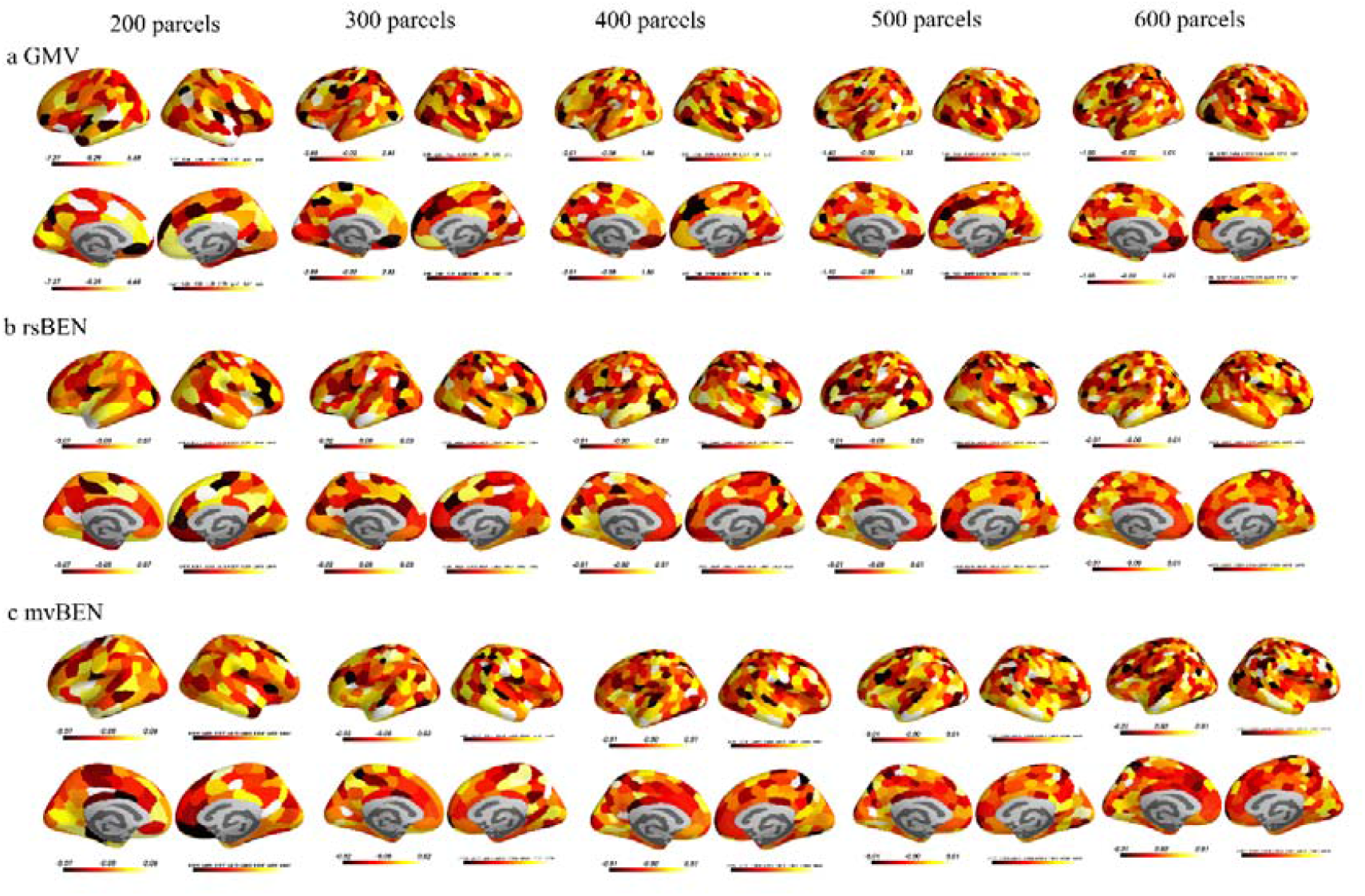
Parcels weights for cognitive scores

**Fig3.**
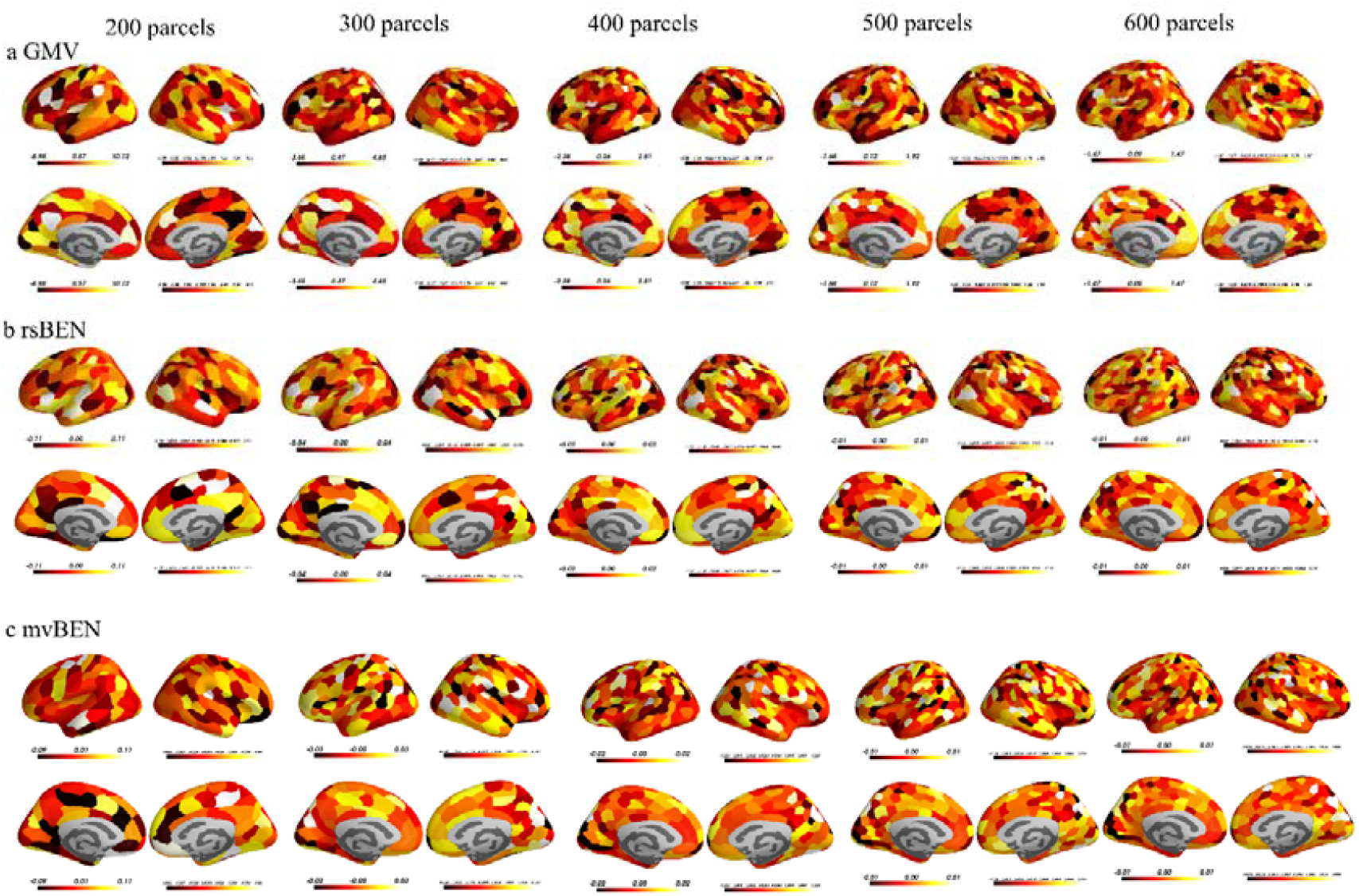
Parcels weights for emotion scores

For predicting cognitive scores, GMV is primarily associated with the left ventromedial prefrontal cortex (VMPFC), the dorsomedial prefrontal cortex (DMPFC), the temporoparietal junction (TPJ), the right occipital lobe, the right motor cortex, and the insula. On the other hand, rsBEN is mainly linked to the left DMPFC, the left anterior temporal lobe (ATL), the right VMPFC, the right insula, the right lateral orbitofrontal cortex (LOFC), and the TPJ. For predicting emotional scores, rsBEN is primarily associated with the left VMPFC, the dorsolateral prefrontal cortex (DLPFC), the left posterior temporal inferior gyrus, the right occipital lobe, the right precuneus, and the right parietal lobe. Meanwhile, mvBEN is mainly linked to the right LOFC, the right insula, the occipital lobe, and the TPJ.

We also calculated the correlations between the prediction coefficients for GMV, rsBEN, and mvBEN using 600 parcels. The results indicated that both rsBEN and mvBEN showed significant correlations in their regression coefficients for predicting cognitive scores (r=0.578, p<0.001) as well as emotion scores (r=0.569, p<0.001). However, there is no significant correlation with GMV. This suggests that rsBEN and mvBEN share similar features, which are not directly related to the features derived from GMV.

## 4. Discussion

Our results demonstrate that prediction models based on BEN are effective in predicting phenotypes. Moreover, BEN-based prediction models outperform GMV-based models, with mvBEN-based models showing better predictive performance compared to rsBEN-based models.

CPM is widely used, and the data from this study have also been used to predict phenotypes by CPM. Compared to CPM we found that BEPM not only outperforms GPM but also demonstrate prediction accuracy comparable to CPM. In Finn’s study (Finn and Bandettini 2021), the highest prediction accuracy for cognitive scores using resting-state data reached r=0.26, while BPM achieved a maximum accuracy of r=0.318. For movie-watching data, the highest prediction accuracy using CPM was r=0.41, whereas the BPM reached r=0.43. For emotional scores, the resting-state CPM showed no significant predictive ability, while BPM achieved a maximum prediction accuracy of r=0.23. Using movie-watching data, the CPM had a prediction accuracy of r=0.18, whereas the BPM reached r=0.293. It is important to note that some issues exist with this comparison, primarily because our results are based on multiple parcels, which may exaggerate our accuracy. Additionally, our selected atlas did not include subcortical regions and the cerebellum, which might have limited the accuracy of the BPM, particularly given the significant roles of the subcortex and the cerebellum in cognition and emotion (Strick, Dum et al. 2009, Forstmann, de Hollander et al. 2017, Pessoa 2017, Schmahmann 2019). Nevertheless, our results indicate that BPM not only surpass GMV-based models but also offer prediction accuracy comparable to CPM, highlighting that BEN, based on local dynamics, can provide extensive information about brain activity.

Future work should explore a wider range of brain atlases to identify the optimal atlas for BPM. The currently selected atlas, generated from connectome, may not align perfectly with the distribution of BEN, particularly at parcellation boundaries. This misalignment could explain why models with more parcels sometimes exhibit lower prediction accuracy compared to models with fewer parcels. In developmental and clinical contexts, accurate prediction models can provide significant practical benefits. Therefore, it is important to investigate the advantages of using BPM in these areas, focusing on their potential benefits for developmental and clinical applications.

**Let me (Donghui Song) continue refining this work after I complete my doctoral thesis defense!**

## Acknowledgements

Data were provided by the Human Connectome Project, WU-Minn Consortium (Principal Investigators: David Van Essen and Kamil Ugurbil; 1U54MH091657) funded by the 16 NI H Institutes and Centers that support the NIH Blueprint for Neuroscience Research; and by th e McDonnell Center for Systems Neuroscience at Washington University.

## Data and code availability

All structural and functional MRI data are available at https://www.humanconnectome.org/

BENtbx is available at https://www.cfn.upenn.edu/zewang/BENtbx.php.

## CRediT authorship contribution statement

Dong-Hui Song: conceptualization, data analysis, visualization, manuscript drafting, and editing. Ze Wang: conceptualization, manuscript editing, supervision, project administration.

